# Communicability distance reveals hidden patterns of Alzheimer disease

**DOI:** 10.1101/2020.04.07.029249

**Authors:** Eufemia Lella, Ernesto Estrada

## Abstract

The communicability distance between pairs of regions in human brain is used as a quantitative proxy for studying Alzheimer disease. Using this distance we obtain the shortest communicability path lengths between different regions of brain networks from Alzheimer diseased (AD) patients and healthy cohorts (HC). We show that the shortest communicability path length is significantly better than the shortest topological path length in distinguishing AD patients from HC. Based on this approach we identify 399 pairs of brain regions for which there are very significant changes in the shortest communicability path length after AD appears. We find that 42% of these regions interconnect both brain hemispheres, 28% connect regions inside the left hemisphere only and 20% affects vermis connection with brain hemispheres. These findings clearly agree with the disconnection syndrome hypothesis of Alzheimer disease. Finally, we show that in 76.9% damaged brain regions the shortest communicability path length drops in AD in relation to HC. This counterintuitive finding indicates that AD transforms the brain network into a more efficient system from the perspective of the transmission of the disease, because it drops the circulability of the disease factor around the brain regions in relation to its transmissibility to other regions.

## 1 Introduction

The human brain is arguably the most complex of all complex systems. At the most basic structural level of interest for neurosciences, the human brain consists of 10^11^ neurons and 10^12^ glial cells, which communicate through neural projections [1]. These cells are then packed into local circuits [2] or large gyri, which define anatomical and functional regions in the brain. The human brain is considered to be outstanding among mammalian brains, it is the largest-than-expected from body size, and it has an overdeveloped cerebral cortex representing over 80% of brain mass [1, 3]. Most of the complexity of these different size-scales of the human brain comes not only from the number of its components, but mainly from the intricate webs of connections linking these components. The emerging field of *network neuroscience* studies the structural and dynamical properties of these webs observed at different size-scales from a variety of noninvasive neuroimaging techniques [4].

The term *pathoconnectomics* has been coined by Rubinov and Bullmore [5] to describe the use of network neuroscience techniques on the analysis of abnormal brain networks (see also [6]). The goals of pathoconnectomics are not only of practical relevance as in the early diagnosis of psychiatric and developmental disorders, stroke, severe brain injury and neurodegenerative diseases, but also in the understanding of their causal mechanisms as pointed out by Raj and Powell [7]. Due to its societal challenge, Alzheimer’s disease (AD) has become a major focus of pathoconnectomic research agenda. AD is the most common neurodegenerative disorder and it represents a major growing health problem for elderly population [8, 9, 10, 11]. It is characterized by a continuous degradation of the patient, which starts with a preclinical stage, a phase of mild cognitive impairment (MCI), and finishing with dementia. The molecular basis of these different stages appear to be linked to the presence of *β*-amyloid (A*β*) in senile plaques and cerebral amyloid angiopathy, as well as tau proteins (tau) in neurofibrillary tangles [12, 13, 14]. For instance, the cognitive decline in AD correlates with tauopathy [15, 16, 17], while the aggregation of A*β* appears to be critical in the early stages that trigger events conducting to tauopathy, neuronal dysfunction, and dementia [18]. Then, it is plausible that these proteins, A*β* and tau, originate in a particular region of the brain and then propagate through neural fibers in a prion-like manner [19, 20, 21, 22, 23].

The hypothesis of the self-propagation of AD in combination with network neurosciences has triggered the use of epidemiological models on networks to simulate the propagation of a disease factor as AD progresses. In particular, Peraza et al. [24] have proposed the use of the Susceptible-Infected (SI) model on networks (see for instance [25, 26, 27]), in which nodes are in two possible states, infected (I) or susceptible (S). The first correspond to brain regions wherein the disease factor is present with high probability, while the second are those free of the disease factor but that can be infected from any infected nearest neighbor. Similar principles have guided Iturria-Medina et al. [28] in modeling the progression of AD under the Network Diffusion Model of disease progression in dementia [29].

Here we start by adopting the SI-model for the propagation of a disease factor in AD. However, we use this model to connect with the theory of network communicability, which has been widely used in network neurosciences (for some applications of communicability in pathoconnectomics see [30, 31, 32, 33, 34, 35, 36, 37, 38, 39, 40]). That is, we will provide a theoretical connection between the network communicability and the probability of a disease factor of propagating from one node to another in a network. Using this connection we will consider a measure that account for the difference between the circulability of this disease factor around a given pair of nodes and its transmissibility from one region to the other. This measure is a Euclidean distance metric–communicability distance–for the corresponding pair of nodes in the network. We then find the length or the shortest communicability paths between every pair of regions in human brains for cohorts of healthy (HC) and Alzheimer diseased (AD) individuals after appropriate normalization. We report in this work that: (i) the shortest communicability path length is orders of magnitude more significant in distinguishing AD from HC than the shortest topological path length; (ii) there is a set of 399 pairs of regions for which there are very significant changes in the shortest communicability path length after AD, (iii) 42% of these significant pairs of brain regions interconnect both brain hemispheres, while 28% connect regions inside the left hemisphere only, in agreement with findings related to the disconnection syndrome. Additionally, 20% of these pairs of affected regions are connecting the vermis with any of the two brain hemispheres, in agreement with recent results; (iv) for 76.9% of these pairs of damaged brain regions there is an increase in the average cliquishness of the intermediate regions which connect them, which implies a significantly higher energy consumption for communication between these regions in AD than in HC.

## 2 Theoretical approach

Here we will use indistinguishably the terms graph and network and follow the classical notation in network theory (see for instance [41]). A *graph G* = (*V, E*) is defined by a set of *n* nodes (vertices) *V* and a set of *m* edges *E* = {(*u, v*) | *u, v* ∈ *V*} between the nodes. The *degree* of a vertex is the number of edges incident to it. A *walk* of length *k* in *G* is a set of nodes *i*_1_, *i*_2_,…, *i*_*k*_, *i*_*k*+1_ such that for all 1 ≤ *l* ≤ *k*, (*i*_*l*_, *i*_*l*+1_) ∈ *E*. A *closed walk* is a walk for which *i*_1_ = *i*_*k*+1_. A *path* is a walk with no repeated nodes. The length of a path is the number of edges in that path. The shortest of all paths connecting the same pair of vertices is known as the shortest topological path. A graph is *connected* if there is a path connecting every pair of nodes. Here we will only consider connected graphs.

Let *A* be the adjacency matrix of the graph, which for simple finite graphs is symmetric, and thus its eigenvalues are real. We label the eigenvalues of *A* in non-increasing order: *λ*_1_ ≥ *λ*_2_ ≥ … ≥ *λ*_*n*_. Since *A* is a real-valued, symmetric matrix, we can decompose *A* into *A* = *U* Λ*U* ^*T*^, where Λ is a diagonal matrix containing the eigenvalues of *A* and 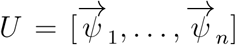 is orthogonal, where 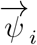 is an eigenvector associated with *λ*_*i*_. Because the graphs considered here are connected, *A* is irreducible and from the Perron-Frobenius theorem we can deduce that *λ*_1_ > *λ*_2_ and that the leading eigenvector 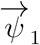, which will be sometimes referred to as the *Perron vector*, can be chosen such that its components 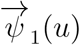 are positive for all *u* ∈ *V*.

### 2.1 Susceptible-infected model

We start by considering a susceptible-infected model of propagation of a disease factor as AD progresses. In this case the brain regions, represented as nodes of the graph, can be in two possible states, infected or susceptible. Susceptible brain regions are those which are free of the disease factor but which are susceptible to get infected from other regions. The infected ones are those in which the probability of disease factor is greater than zero. Let *i* be a node of the graph *G* = (*V, E*) and let *x*_*i*_(*t*) be the probability that node *i* get infected at time *t* from any infected nearest neighbor. If the infection rate is given by *γ* we have [25, 26, 27],

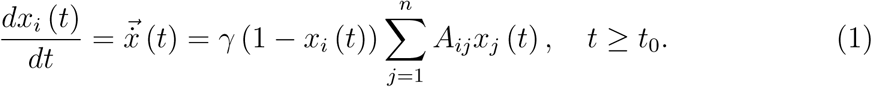

It can be seen that the linearized SI model, namely 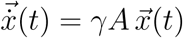 represents an upper bound for the exact SI model, i.e.,

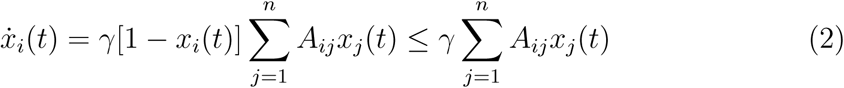

which in matrix-vector for is given by

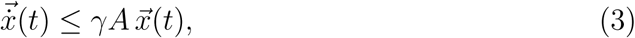

with initial condition 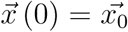, ∀*i* and ∀*t*. The solution of the linearized SI model 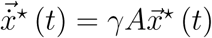 is given by:

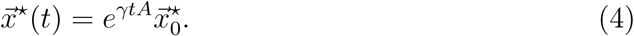

However, the solution of the linearized SI model has two important problems. The first is that the semigroup (*e*^*tA*^)_*t*≥0_ is unbounded, i.e., lim_*t→∞*_ *‖ e*^*tA*^‖ = *∞*, which poses a major problem for the use of this linearized model as a model of the SI propagation scheme. The reason is that *x*(*t*) is a probability and as such it has to be bounded as 0 ≤ *x*(*t*) ≤1. The second is that the solution (4) is a good approximation to the solution of the SI model only for 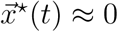, which makes this solution useless for following the propagation of the AD.

To sort out this problem we will follow here Lee et al. [42], who proposed the following change of variable to avoid the aforementioned problems with the solution of the linearized SI model:

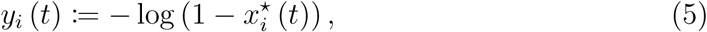

which is an increasing convex function. Then, as 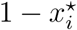 is the probability that node *i* is not infected at a given time, the new variable *y*_*i*_ (*t*) can be interpreted as the information content of the node *i* or surprise of not being infected (see, e.g., [43]). Let us then suppose that at *t* = 0 the probability that every region of the brain gets infected is the same, i.e., at the beginning every node has the same probability *β* to be infected and to be the one from which the disease propagates. That is,

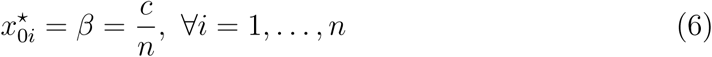

In this case, the solution of the upper bound of the SI model is:

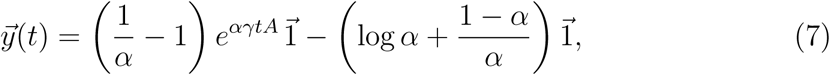

where *α* = 1 – *β* is a constant. Also, if we fix *γ* and *t* we have that this solution can be written as

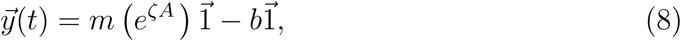

for constants *m,ζ* and *b*. Here the constant *ζ* groups the previous parameters *γ, α* at a given time t. We will call 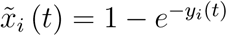 the approximate solution of the SI model, which is always bounded between zero and one as needed.

In order to see the differences between the exact solution of the SI model *x*_*i*_(*t*), the linearized ones 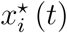 and the approximate solution after the change of variable 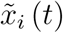, we plot the progression of the number of infected nodes in an Erdős-Rényi random network with *n* = 100 and connection probability *p* = 0.1. We simulate the progression of the disease starting the infection with a fraction of 0.01 infected nodes in the network. The results are illustrated in Figure 1 for infectivity rates *β* = 0.001 (a) and *β* = 0.002 (b). As can be seen the lineralized model (dotted black line) is a bad approximation to the exact solution (broken blue line) as it quickly diverges. However, the approximate solution obtained by the change of variable (solid red line) is a tight upper bound for the exact solution, and it will be used here for further analysis.

**Figure 1:**
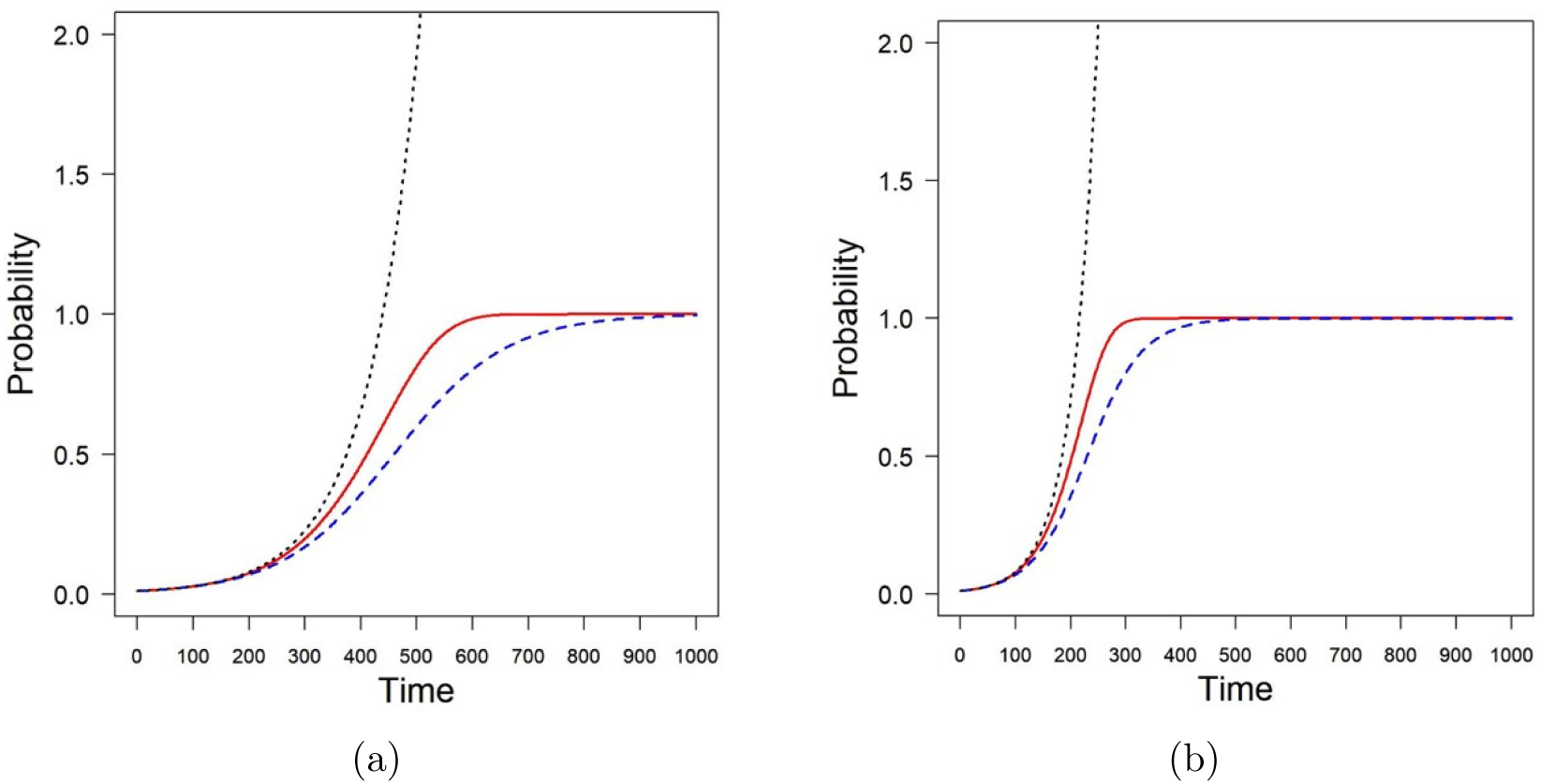
Simulation of the progression of a disease with the SI model (broken blue line) in an Erdős-Rényi random network with *n* = 100 and connection probability *p* = 0.1. The lineralized solution is represented by dotted black line and the approximate solution using the change of variables is represented as a solid red line. The panels correspond to infectivity rates *β* = 0.001 (a) and *β* = 0.002 (b).

### 2.2 The communicability connection

It is clear from Eq. (8) that the solution 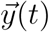 of the upper bound of the SI model depends linearly of 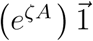. This term is the sum of the corresponding rows of the exponential of the adjacency matrix. For an individual node *i* it is known as the total communicability of the corresponding node [44] and it can be written as

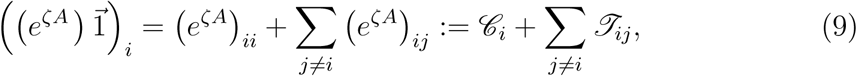

where the first term in the right-hand side is the subgraph centrality [45] of the node and the second one is the sum of the communicability functions [46] from the node *i* to the rest of the nodes of the network (see also [47]). In terms of the propagation of a disease factor, *ℒ*_*i*_ represents the circulability of the disease factor around the node *i*. The second term represents the transmissibility from/to the node *i* to/from the rest of the nodes of the network. If we concentrate on the effect of the node *i* on another node *j*, then *ℒ*_*i*_ represents the capacity of the node *i* of increasing the probability of infesting itself and *𝒯*_*ij*_ is the capacity of *i* of infestating *j*. Thus, because node *j* is doing the same, the term

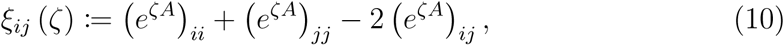

represents the difference between the capacities of both nodes of increasing the probability of infesting themself to that of infesting each other. A large value of *ξ*_*ij*_ (*ζ*) indicates that the disease factor gets trapped circulating at the nodes *i* and *j*, which form two islands with little transmissibility among them. A small value, however, indicates that such transmissibility is relatively large in relation to the internal circulability at the nodes, i.e., the nodes have a bridge between them. We consider that this measure is important for the study of Alzheimer disease because it should allow us to investigate whether the disease produces a patchy environment of brain regions which form islands with little transmissibility among them. The function *ξ*_*ij*_ (*ζ*) can be written as 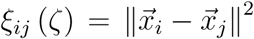, where 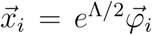 with 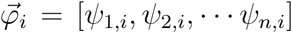 where *ψ*_*k,i*_,is the *i*th entry of the *k*th eigenvector associated with the eigenvalue *λ*_*k*_ of *A*. Consequently, *ξ*_*ij*_ (*ζ*) is a Euclidean distance between the nodes *i* and *j* in the network. We call it the *communicability distance* between the two nodes [48, 49]. The vector 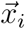 is the position vector of the node *i* in a Euclidean hypersphere of dimension *n* [50, 51].

### 2.3 Shortest communicability paths

The communicability distance *ξ*_*ij*_ (*ζ*) can be calculated for any pair of nodes (connected or not) in the graph. Thus, we can obtain a communicability distance matrix [48]

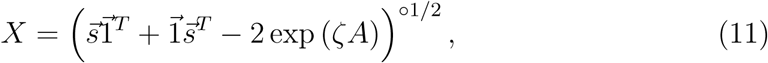

where 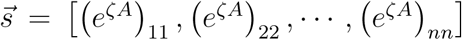 is a vector whose entries are the main diagonal entries of the corresponding matrix function, 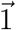 is an all-ones vector and º indicates an entrywise operation. However, we assume here that in a network “information” flows through the edges of the graph, such that it uses certain paths connecting the corresponding pair of nodes. In order to find the shortest communicability paths between two nodes we proceed as follows. We construct the communicability-weighted adjacency matrix of the network [52]:

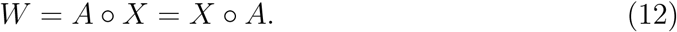

Then, the shortest communicability path between two nodes is the shortest weighted path in *W*. That is, the shortest communicability path between two nodes *i* and *j* for a given *ζ* > 0 is the path that minimizes the communicability distance between every pair of nodes in the corresponding path. We have proved analytically that when *ζ →* 0 the shortest communicability path between any pair of nodes *i* and *j* is identical to the shortest (topological) path between the two nodes [53]. That is, the shortest (topological) path is a special case of the shortest communicability path in a network. In this work we will consider the case *ζ ≡* 1 which we will call “shortest communicability path” and the case *ζ →* 0 which we will call “shortest topological path”. Notice that the length of the shortest communicability path is the sum of the weights (communicability distances) for the edges in that path. For an example see Fig. 2. Here we will keep *ζ* = 1 due to the lack of any experimental value that can guide us for selecting a more appropriate value. Also, we should have in mind that decreasing the values of this parameter close to zero will make the shortest communicability paths to look very similar to shortest paths, while increasing it over unity will make these paths very long indeed. Thus, we left for a further work the analysis of the influence of this parameter in the study of AD.

**Figure 2:**
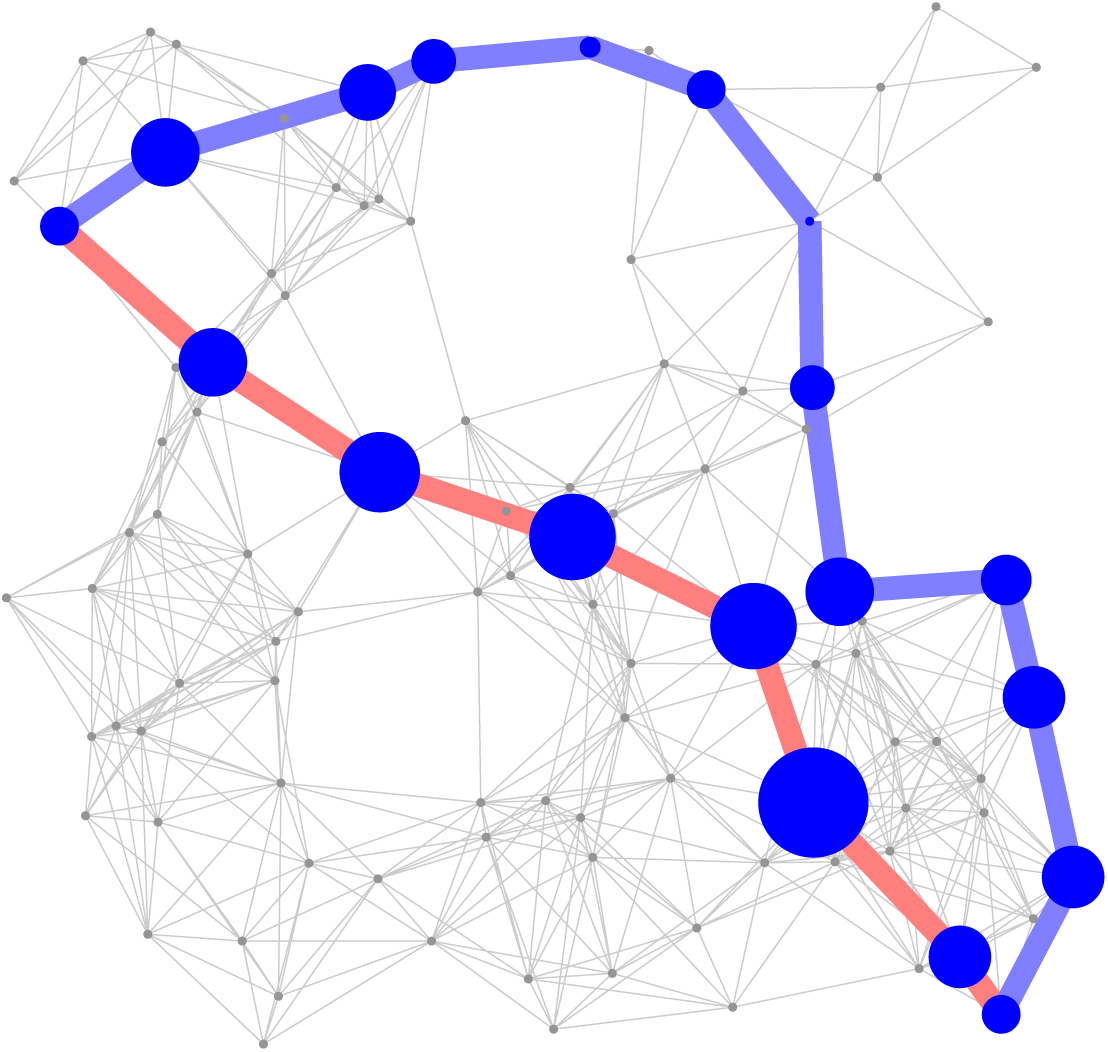
Illustration of the shortest communicability path (blue thick lines) and the shortest topological path (red thick lines) between a pair of nodes in a random geometric network. The nodes in both shortest paths are highlighted with blue color and with sizes proportional to their degrees. The rest of the nodes are in gray color and with a fixed size. The shortest topological path goes through nodes averaging node degree equal to 14. The shortest communicability path goes through nodes averaging degree 8.3. Additionally the average subgraph centrality of the nodes in both paths are 18837.3 (shortest topological path) and 4636.6 (shortest communicability path), respectively.

## 3 Dataset and image processing

The data set used consists of DWI scans and anatomical T1 scans of 88 subjects, 48 Healthy Controls (HC) and 40 Alzheimer’s Disease (AD) patients from the publicly available ADNI database. After preprocessing the images, a tractography pipeline was implemented by using the MRtrix software library. The main steps of the whole processing, which are well-established in the literature, are shown in Fig. 3.

**Figure 3:**
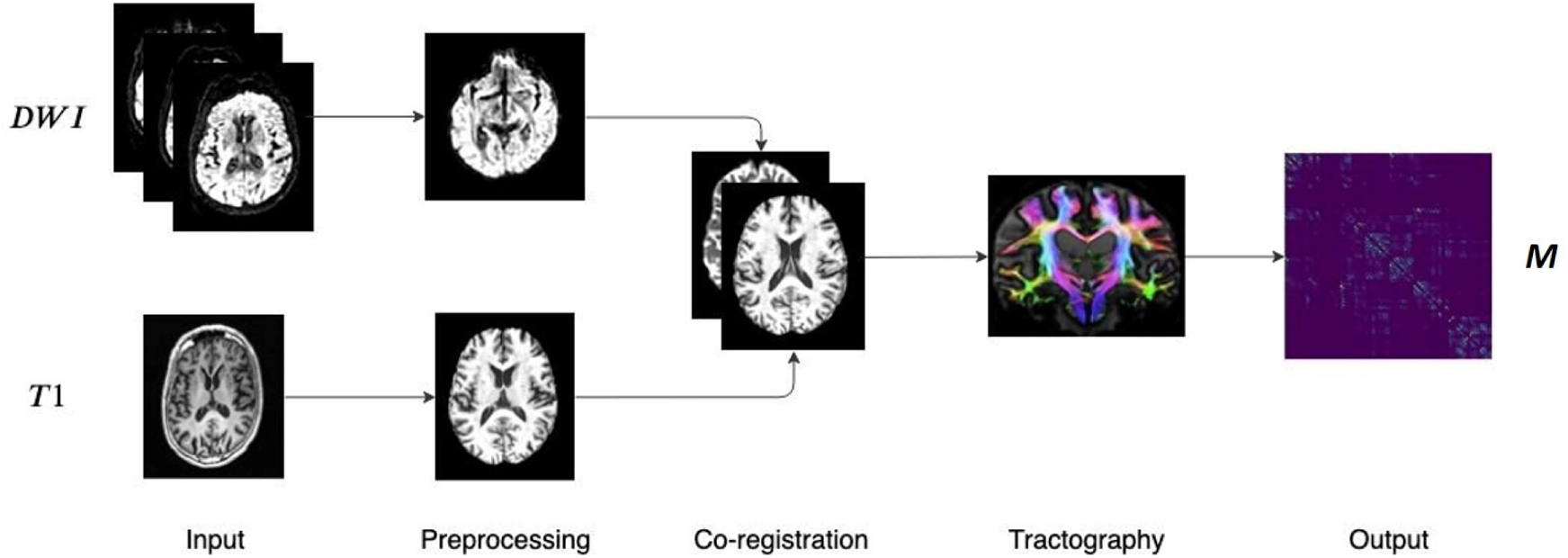
Main steps of the image processing pipeline. DWI and T1 weighted scans are preprocessed and co-registered. Then, after the fiber orientation distribution estimation, probabilistic tractography is performed resulting in a weighted connectivity matrix.

First, a denoising step was performed in order to enhance the signal-to-noise ratio of the diffusion weighted MR signals in order for reducing the thermal noise due to the stochastic thermal motion of the water molecules and their interaction with the surrounding micro-structure [54]. Head motion and eddy current distortions were corrected by aligning the DWI images of each subject to the average *b*_0_ image. The *brain extraction tool* (BET) was then used for the skull-stripping of the brain [55]. The bias-field correction was used for correcting all DWI volumes. The T1 weighted scans were preprocessed by performing the standard steps: reorientation to the standard image MNI152, automatic cropping, bias-field correction, registration to the linear and non-linear standard space, brain extraction. The following step was the inter-modal registration of the diffusion weighted and T1 weighted image.

After the preprocessing and co-registration steps, the structural connectome generation was performed. First, we generated a tissue-segmented image tailored to the anatomically constrained tractography [56]. Then, we performed an unsupervised estimation of WM, gray matter and cerebro-spinal fluid. In the next step, the fiber orientation distributions for spherical deconvolution was estimated [57]. Then a probabilistic tractography [58] was performed by using dynamic seeding [59] and anatomically-constrained tractography [60], which improves the tractography reconstruction by using anatomical information by means of a dynamic thresholding strategy. We applied the spherical-deconvolution informed filtering of tractograms (SIFT2) methodology [59], providing more biologically meaningful estimates of the structural connection density and a more efficient solution to the streamlines connectivity quantification problem. The obtained streamlines were mapped through a T1 parcellation scheme by using the AAL2 atlas [61], which is a revised version of the automated anatomical atlas (AAL) including 120 regions. Finally, a robust structural connectome construction was performed for generating the connectivity matrices [62]. The pipeline here described has been used in recent structural connectivity studies, for example [63, 64, 65, 66]. The output was a weighted connectivity matrix for each subject. Out of these 120 nodes, 24 were removed from all networks, in order to obtain only graphs without isolated nodes. Finally, a 96 × 96 matrix M for each subject was obtained.

All matrices were binarized by considering only the edges with *m*_*ij*_ > 0 and the adjacency matrix *A* was obtained. The communicability distance matrix was calculated for each binary matrix and it was multiplied by the adjacency matrix *A* obtaining a weighted matrix *W*. A shortest path algorithm was performed on this matrix thus obtaining a matrix whose entries represent the shortest communicability paths between node pairs in the network. Starting from the adjacency matrix *A*, the shortest path length matrix, whose entries represent the shortest paths between node pairs, was also calculated. A group-wise statistical analysis was performed in order to find brain region pairs with statistically significant difference between HC and AD in shortest path length and shortest communicability path length. In order to make the statistical analysis more robust, permutation tests were performed by randomly assigning subjects to the two comparison groups 1,000 times. Differences were considered significant if they did not belong to 95% of the null distribution derived from the permutation tests (corrected *p*-value < 0.05). The False Discovery Rate (FDR) was used for multiple comparison correction. The same study could also be done by considering the weighted matrix but we follow here the more traditional approach on binary matrix. The weighted case could be addressed in future work.

## 4 Statistical analysis

### 4.1 Sensitivity analysis

Our first task here is to analyze the sensitivity of the shortest communicability and the shortest topological path lengths to detect significant changes in the brain connectivity after Alzheimer disease. For that purpose we proceed as follow. For each connectivity matrix we calculate both the shortest communicability path length and the shortest topological path length matrices. In this case we use permutation tests as the statistical significance analysis. For instance, let us consider the nodes *v*_*i*_ and *v*_*j*_. We then calculate the length of the communicability shortest path between these nodes for each of the healthy 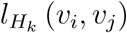 and diseased individuals 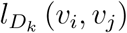 and then obtain the respective average values, 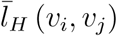 and 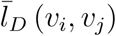. Using these values we obtain 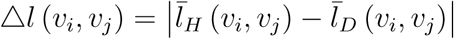. Now we proceed to a randomization of each individual into the two classes, i.e., healthy and diseased, obtaining 1,000 subsets of random HC and 1,000 subsets of random AD. We then calculate 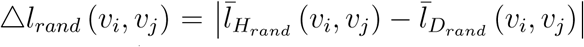, where 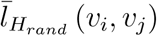 and 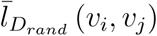 are computed as before but using the random sets of healthy and diseased individuals, respectively. Finally, we compare the null distribution of Δ*l*_*rand*_ (*v*_*i*_, *v*_*j*_) with the true value Δ*l* (*v*_*i*_, *v*_*j*_). Therefore we conclude that Δ*l* (*v*_*i*_, *v*_*j*_) is significant if it does not belong to 95% of the null distribution, which is carried out by calculating the *p*-value of the permutation test. Here we consider significant the nodes with corrected *p*-value < 0.05. We use both False Discovery Rate (FDR) and Bonferroni correction for the multiple comparisons correction. We do these calculations for the shortest communicability path as well as for the shortest topological path.

### 4.2 Effects of threshold selection and normalization

Here we first consider the effects of the thresholding process on the significance of the results obtained by using the current approach. The brain networks used in this work, as usually in many brain network studies, are constructed by using probabilistic tractography. For this reason weak connections can introduce noisy effects. Therefore, the first thing that we need to investigate is how different thresholds to transform these weighted matrices into binary (adjacency) matrices affect the results. The second important question is related to the comparison of networks with very different topological characteristics to avoid the extraction of trivial facts. That is, it is very plausible that the brain networks of AD patients differ significantly in a few “trivial” topological aspects from those of healthy individuals. For instance, the edge density can change dramatically between HC and AD networks. This may produce the false impression that AD mainly produces a sparsification of the brain network which hides important structural factors produced by the disease. To avoid these problems we will provide a normalization of the communicability geometric parameters used in this study as described below.

First, we will proceed to change the threshold at which the adjacency matrices are generated. We start as usual by calculating the mean matrix for the HC subjects, which results in a weighted matrix whose entries range from 0 to 1. Each entry represents the frequency at which the corresponding edges occur among the HC matrices.

This matrix is then thresholded by varying the threshold *τ* as 0 ≤ *τ* ≤ 0.9 obtaining a binary matrix to be used as a mask. The adjacency matrices of all subjects are then projected onto this mask. This procedure resulted in ten sets of adjacency matrices, one set for each threshold value. We use the same thresholding procedure described in [30], which is also similar to the procedure used for example in [67]. In Fig. 4 we illustrate the results of the statistical significance of the different thresholds studied here for both FDR (a) and Bonferroni correction (b). The first shocking result is the extremely low significance of the shortest topological paths according to both multiple comparisons correction methods for all values of the threshold. According to FDR for almost all values of thresholds the ratio of significant node pairs is more than 35 times higher than that for the shortest topological paths, while according to the most restrictive Bonferroni correction the ratio of significant node pairs is quite dependent on the threshold value and the highest ratio of significant node pairs is obtained for *τ* between 0.5 and 0.9.

**Figure 4:**
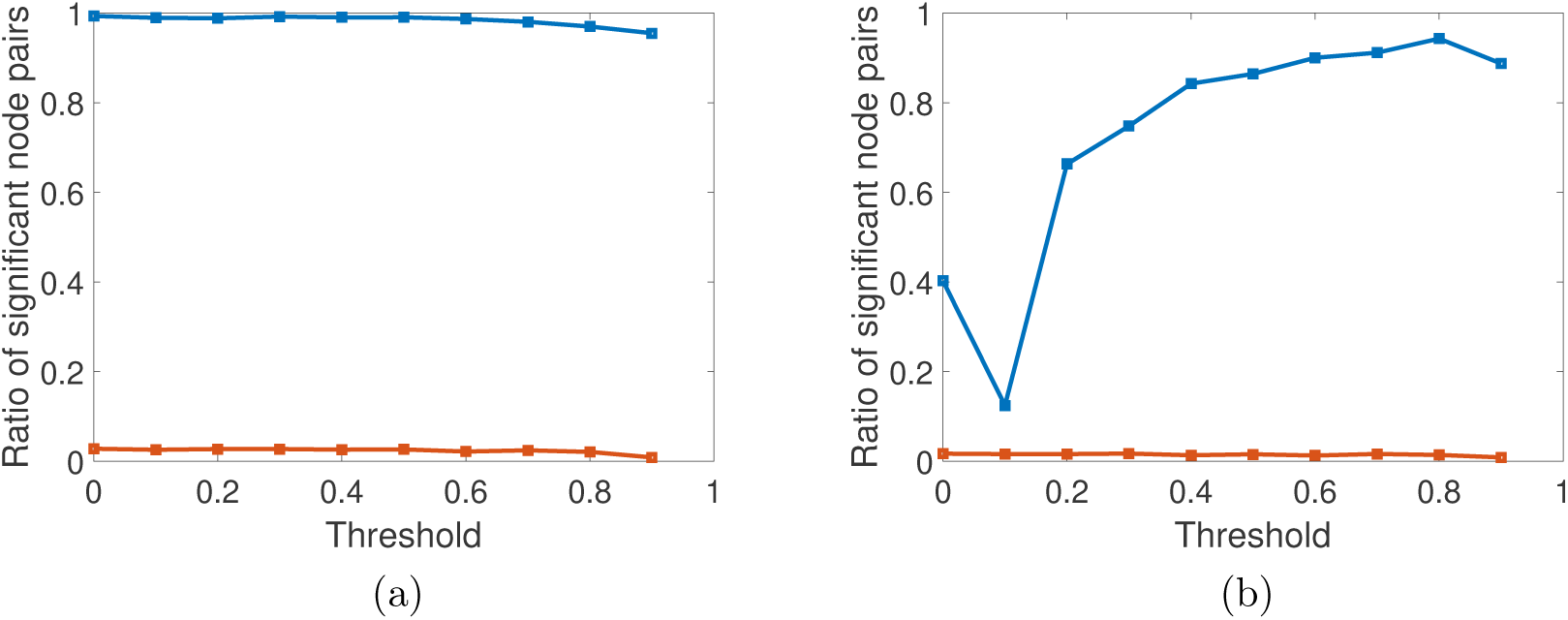
Fraction of node pairs with statistically significant different values of shortest communicability path length (blue squares) in HC and AD compared to the number of node pairs with statistically significant different values of shortest path length (red squares), at different threshold values for FDR (a) and Bonferroni correction (b).

Then, for each value of the threshold we calculate the communicability distance matrix *X* (*P*_*i*_) of subject *i*. We then proceed to normalize such matrix as follow. Let us call *S* (*P*_*i*_) the shortest communicability path length matrix of the subject *P*_*i*_, let *m*(*P*_*i*_) be the number of edges of subject *i*, and let *A*(*P*_*i*_) the adjacency matrix of subject *i*. The average communicability distance of the edges of the network is then calculated for each subject:

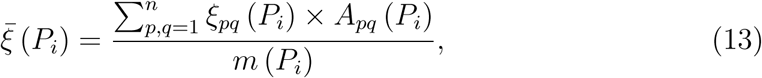

from which we obtain the normalized shortest communicability path length matrix *Ŝ*(*P*_*i*_) as:

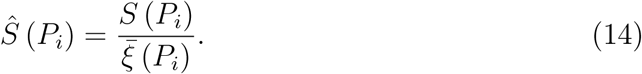

For instance, for the network *G* in Fig. 5 the value of 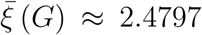, then the normalized shortest communicability path length for the pair labeled as 1, 6 is *Ŝ*_1,6_ (*G*) ≈ 1.939, which corresponds to the path 1 – 9 – 8 – 7 – 6. In contrast, the normalized length of the communicability path 1 – 2 – 4 – 6 is 2.049, which clearly indicates that this is a longer path in term of the communicability distance than the path path 1 – 9 – 8 – 7 – 6. Notice that 1 – 2 – 4 – 6 is the shortest topological path between the nodes 1 and 6.

**Figure 5:**
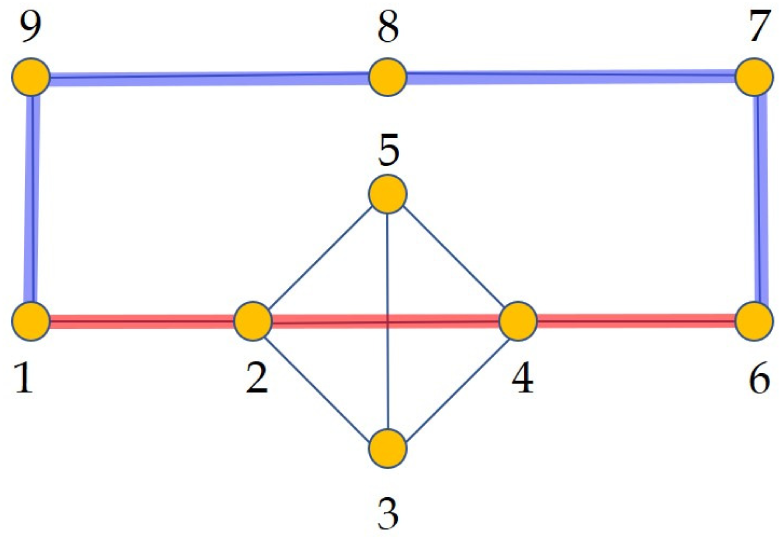
Illustration of a simple graph used to explain the normalized shortest communicability path length. In red we illustrate the shortest path between the nodes 1 and 6 and in blue the shortest communicability path between the same nodes. The normalized communicability shortest path length for the path marked in blue is *Ŝ*_1,6_ (*G*) ≈ 1.939, while for that in red is *Ŝ*_1,6_ (*G*) ≈ 2.049.

We then analyze the significance for the normalized communicability distance matrices, for each threshold studied here in the two cases of the False Discovery Rate (FDR) and the Bonferroni. In Fig. 6 (a) we illustrate the ratio of significant node pairs vs. the threshold for FDR and in (b) the same for the Bonferroni correction. We also provide the same results for the shortest topological paths. The first interesting result is the dramatic difference between the ratios of significant node pairs obtained from the shortest communicability paths and from the topological ones. While for the shortest communicability paths we have very significant ratios for both statistical parameters for certain values of the threshold, for the shortest paths we always observe very low ratio of significant node pairs both for FDR and Bonferroni correction. The second very interesting feature of this analysis is the fact that for both statistical criteria the normalized shortest communicability paths makes a great differentiation of both groups for a threshold of *τ* = 0.5. Notice the nonmonotonic behavior of both statistical criteria versus the threshold, which peak at the before mentioned value.

**Figure 6:**
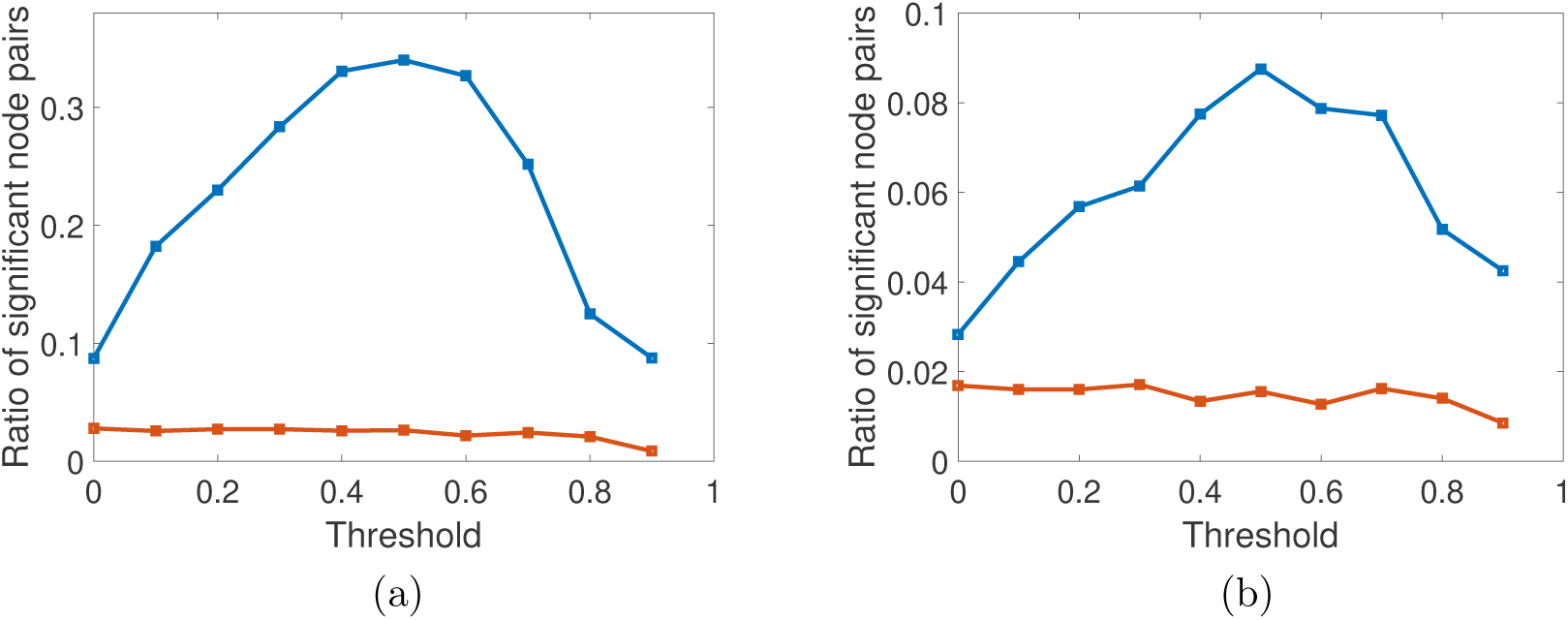
Fraction of node pairs with statistically significant different values of normalized shortest communicability path length (blue squares) in HC and AD compared to the number of node pairs with statistically significant different values of shortest path length (red squares), at different threshold values for FDR (a) and Bonferroni correction (b).

Now we focus only on the results obtained after the normalization of the communicability distance matrix and for the threshold found as providing the best results. In Fig. 7 we illustrate these results using Manhattan plots for the significance of node pairs according to FDR (panels a and b) as well as for Bonferroni correction (panels (c and d). Notice that the horizontal red line corresponds to the significance, i.e., – ln (0.05) for FDR and –ln (0.05*/κ*), where *κ* is the number of comparisons, for Bonferroni. All the points over the red line represent significant node pairs. According to the FDR correction there are 1551 significant node pairs for the normalized communicability distance against 120 ones according to the shortest topological paths.

**Figure 7:**
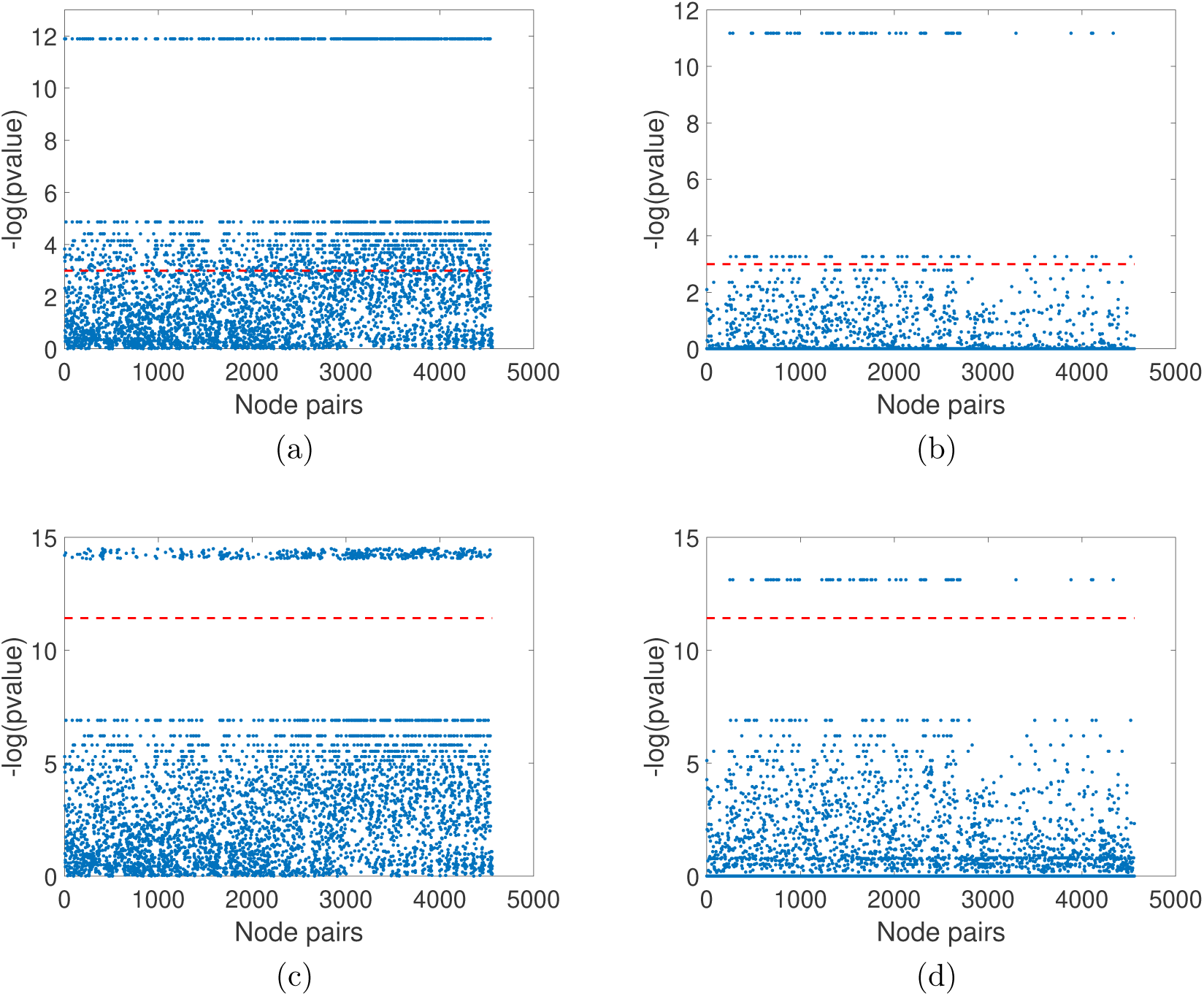
Results of the group-wise statistical analysis using FDR (a and b) and the Bonferroni index (c and d) for the normalized communicability distance matrices with threshold *τ* = 0.5 (a and c) as well as for the shortest topological paths. The node pairs above the red line have significantly different values in HC and AD.

According to the Bonferroni correction there are 399 significant node pairs for the normalized communicability shortest paths against 71 for the topological ones. We should remark that FDR controls the expected proportion of false positives, while Bonferroni controls the overall probability of making at least one false discovery. Then, because the Bonferroni correction is a more restrictive measure we will consider only the 399 significant node pairs identified by this measure. Notice that even with such restrictive criterion the shortest communicability path identifies more than 7 times the number of significant node pairs identified by the topological shortest paths.

In closing, we observe a huge difference in the sensitivity of the communicability shortest paths respect to the shortest topological ones to the change in the brain connectivity produced by Alzheimer disease. In other words, while the length of the shortest topological paths appear almost unaltered after the appearance of AD, the length of the communicability ones is affected in almost all the pairs of brain regions.

Before closing this section we would like to remark the huge differences produced by the normalization procedure used in the current work as a way to restrict our analysis to those highly nontrivial structural features relevant to Alzheimer disease. When there is no normalization in FDR case the shortest communicability path identifies 4,524 node pairs significantly different in the two diagnostic groups, out of the 4,560 possible total node pairs. That is, 99.21% of the node-pairs are affected. Conversely, for the topological shortest path length only identifies 124 node pairs, which represents only 2.72% of node-pairs affected. Without normalization we can also study the influence of the threshold under the ratio of significant node pairs in Bonferroni case. In this case, the best results are obtained for *τ* = 0.8. Then, there are 4,300 node pairs significantly different in the two diagnostic groups according to the shortest communicability path, and 95 according to the shortest topological ones. These values represent 95.61% and 2.08% node-pairs affected, respectively. These results show the extraordinary value of the normalization criterion used in reducing the number of significant node pairs to a handful set of highly significant ones.

## 5 Discovering structural patterns of Alzheimer disease

Considering this threshold value, the average shortest communicability path length matrix was calculated for HC and for AD. Then, we have obtained the difference between the average matrices for the HC minus that of AD (Fig 8 (a)). For the sake of comparison we also obtained such differences for the shortest path lengths between every pair of nodes (Fig 8 (b)). We used a divergent colormap centred at zero to represent these differences and the differences for the shortest topological path length were set in the same scale of the differences of the normalized shortest communicability path length. It is interesting to note how this representation allows to visualize a different distribution of colours in the two heatmaps. In particular if we consider the histogram of the values of one of the rows of the heatmaps (the row corrisponding to node 35 is considered as an example), the histogram derived from Fig 8 (a) is a bimodal distribution (Fig 8 (c)), while the one derived from Fig 8 (b) is a skewed distribution centered at zero (Fig 8 (d)). Moreover looking at Fig 8 (a), two different behaviors of the distribution of values of a single row can be observed. In particular for some rows the average difference, between HC and AD, of normalized shortest communicability path length with the other nodes of the network is mostly positive, while for other nodes it is mostly negative. Thus the nodes seems to be clustered according to this different behavior. In order to show how these two different groups are distributed in the brain, we have represented the brain regions on a glass brain coloring the corresponding nodes according to the median of the distribution of the values over row (Fig. 9). Also the dimension of the nodes is descriptive of the median value. The cluster of nodes with the highest median values includes Cerebellum, Vermis and Amygdala.

**Figure 8:**
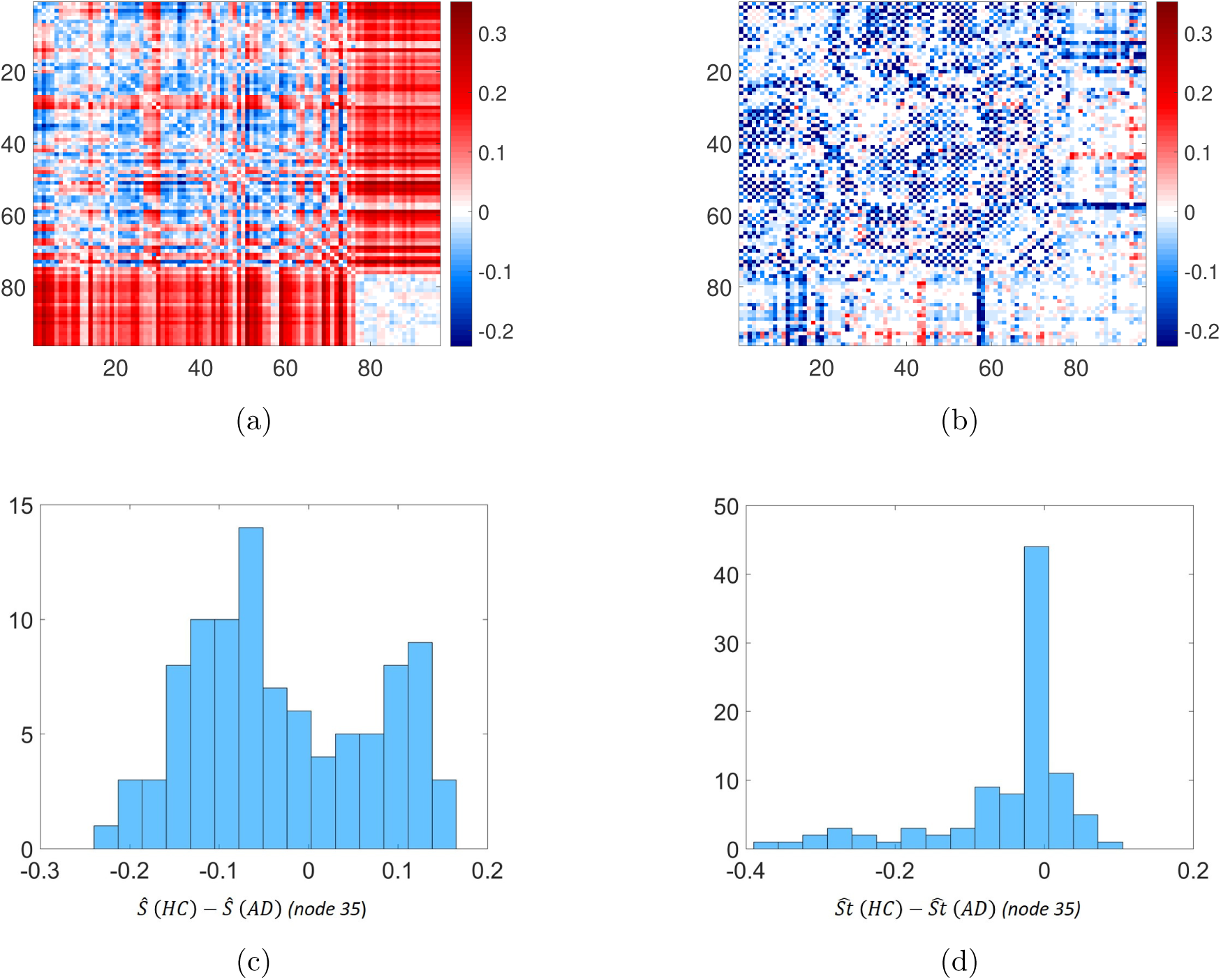
(a) Heatmap of the difference between the averaged normalized shortest communicability path length for HC and AD. (b) Heatmap of the difference between the averaged shortest topological paths length for HC and AD, in the same scale of (a). (c) Distribution of the values of one row (row 35 is considered as an example) of heatmap (a). (d) Distribution of the values of one row (row 35 is considered as an example) of heatmap (b).

**Figure 9:**
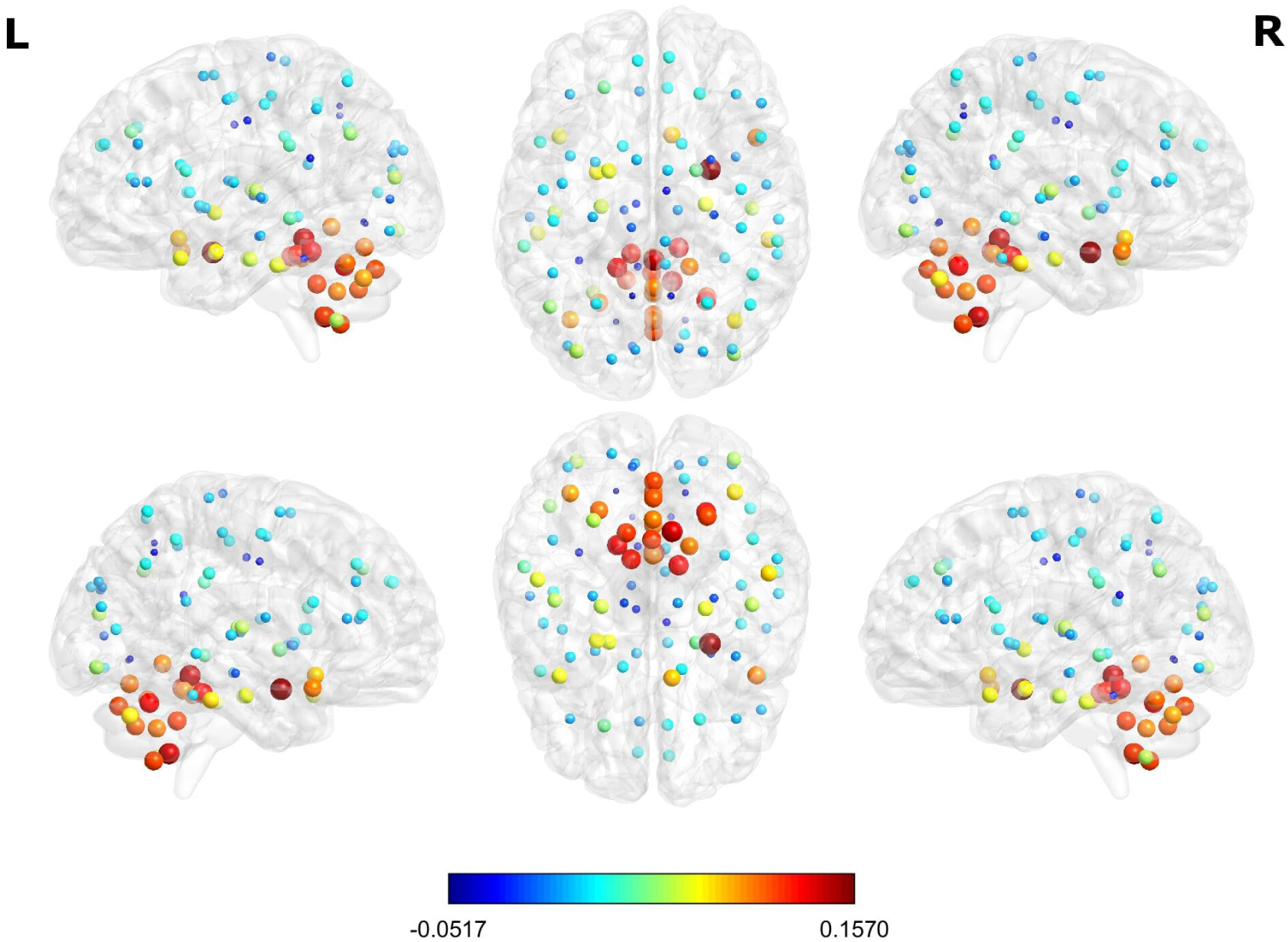
The glass brain shows for each node the median of the distribution of the difference in HC and AD of the mean normalized shortest communicability path length with all the other nodes of the network; the node colour and dimension are descriptive of these values. The different views shows the lateral and medial sides of each hemisphere, and the dorsal and ventral side.

From these heatmaps we can observe that there are pairs of nodes for which AD increases the shortest communicability and topological paths while for others it decreases them. The difference between the distributions of *Ŝ* (*HC*) and *Ŝ* (*AD*), i.e., when *P*_*i*_ is a healthy or Alzheimer diseased individual, respectively, is statistically significant.

A group wise statistical analysis using permutation tests with multiple comparison correction (both False Discovery Rate and Bonferroni correction) was performed in order to find which node pairs have a significantly different value of the normalized shortest communicability path length in HC and AD. This can allow to restrict the focus only on these node pairs, among all the possible node pairs. This procedure was applied for all thresholds.

Let us call *Δ*_*ij*_ = *Ŝ* _*ij*_ (*HC*) – *Ŝ*_*ij*_ (*AD*) the difference of the average normalized shortest path communicability distance for the edge (*i, j*) in both the healthy and the Alzheimer diseased cohorts. Let us then call 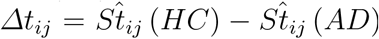 the difference of the average shortest topological path length. Then, we can observe that among the node pairs with statistically significant different values of the average normalized shortest communicability path length, there are both node pairs with *Δ* > 0 and *Δ* < 0. Instead for the node pairs with statistically significant different values of the average shortest topological path length it is always *Δ*_*St*_ < 0. For example, Fig. 10 shows the histograms of *Δ* and *Δ*_*St*_ (we generally call the difference of an average shortest path measure) for the significant node pairs for the best threshold value (*τ* = 0.5).

**Figure 10:**
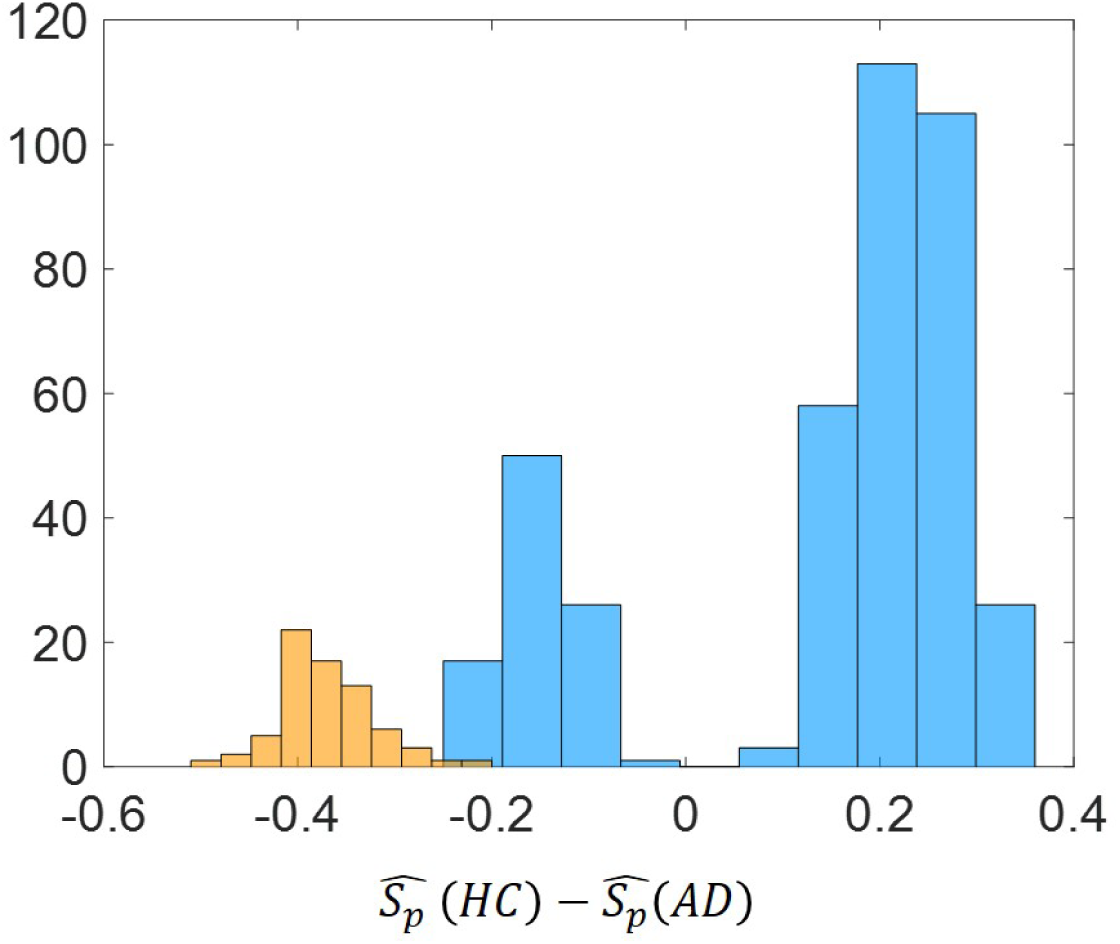
Histogram of *Δ*_*Sp*_ = *Ŝp* (*HC*) − *Ŝp* (*AD*) for the significant node pairs for the best threshold value (*τ* = 0.5). Blue color refers to the difference of the average normalized shortest path communicability distance while orange color refers to difference of the average shortest topological path length.

The values of *Δ*_*ij*_ for the set of 399 node pairs with significantly different average normalized shortest communicability path length for the best threshold (*τ* = 0.5) which were selected according to the Bonferroni correction are illustrated in Figure 11.

**Figure 11:**
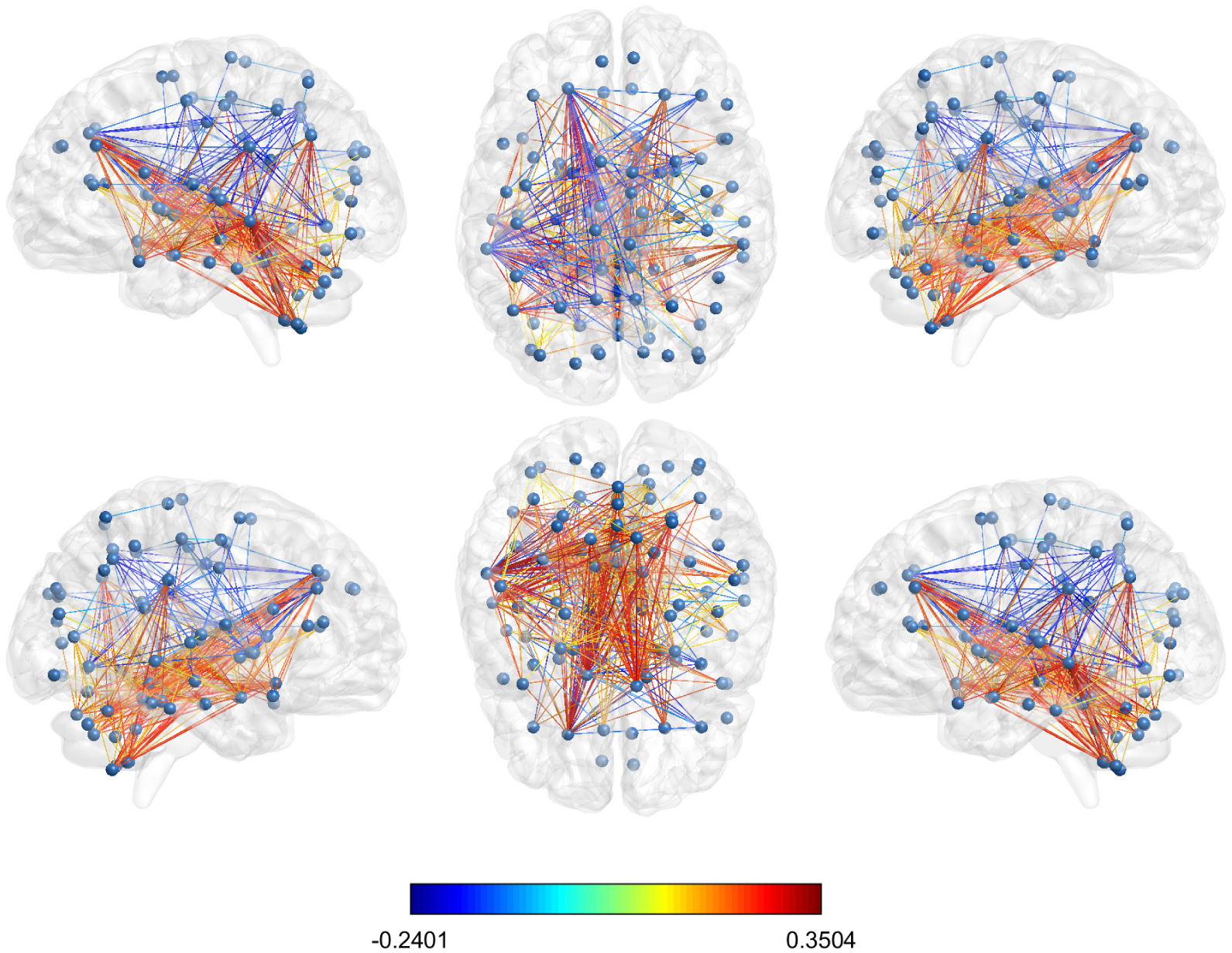
Glass brain visualization of the difference between the mean normalized shortest communicability path length of the significant edges in HC and AD; the edge colour is descriptive of these values. The different views shows the lateral and medial sides of each hemisphere, and the dorsal and ventral side.

A deeper analysis of these node pairs provide some illuminating information about the structural influence of Alzheimer disease. Let us resume this information as follow. From the 399 significant node pairs considered here:

1. 110 (27.6% of significant pairs) connect regions in the left hemisphere;
2. 41 (10.3% of significant pairs) connect regions in the right hemisphere;
3. 167 (41.8 of significant pairs) connect one region of the left with one region of the right hemisphere;
4. 31 (7.8% of significant pairs) connect the Vermis to the right hemisphere;
5. 50 (12.5% of significant pairs) connect the Vermis to the left hemisphere.

These results indicate that almost one half of all the pairs of nodes for which there is a significant difference in the shortest communicability paths after Alzheimer disease connect both brain hemispheres. This result supports the disconnection hypothesis of this disease in which the damage produced by AD can be attributed not only to specific cerebral dysfunctions but also to disconnection processes between different cerebral areas [68, 69]. In particular, the disconnection between inter-hemispheric regions have been widely discussed in the literature and reviewed by Delbeuck et al. [70]. They compiled evidences about the hypothesis of the AD as a disconnection syndrome from neuropathological data, the electrophysiological and neuroimaging data as well as from neuropsychological data. Some of the earliest evidence supporting this hypothesis point out to the fact that there is a disorganized functional activity between the two hemispheres in the early stages of AD and a loss of positive correlations between the hemispheres, which suggest a breakdown of the interhemispheric functional association. Other evidences reviewed by Delbeuck et al. [70] are a decrease of associative white matter fibers in the corpus callosum splenium of AD patients, the existence of a modification in the functional interactions between the hemispheres, and the existence of lower coherence between the hemispheres in mild to moderate AD patients compared to controls, suggesting a disturbance of the interhemispheric functional connectivity in AD. More recently, Wang et al. [71] obtained experimental results which demonstrate that there are “specific patterns of interhemispheric functional connectivity changes in the AD and MCI, which can be significantly correlated with the integrity changes in the midline white matter structures”. And Qiu et al. [72] reported homotopic inter-hemispheric functional connectivity disruption in AD but not MCI.

Another third of these significant edges are located inside the left hemisphere. These results parallels those supporting the hypothesis that AD evolve first, faster, and more severely in the left hemisphere than in the right [73, 74]. As shown by Thompson et al. [73], the spreading wave of gray matter loss were asymmetric in both hemispheres, with the left one having significantly larger deficits and with faster local gray matter loss rates than the right one. Their finding also correlated with progressively declining cognitive status. Finally, we have found 81 pairs which connect the vermis either with the right or the left hemisphere. Recently the role of cerebellar gray matter atrophy in Alzheimer’s disease has been studied [75, 76]. Jacobs et al. [75] have found that in the early stages of AD the vermis and posterior lobe of the cerebellum are affected, also confirming previous results by Mavroudis et al. [76] who reported severe damage in the Purkinje cells from the vermis of the cerebellum in 5 AD patients.

### 5.1 Transmissibility and circulability of Alzheimer disease factor

As explained before, among the pairs of nodes which have significant differences between healthy and Alzheimer diseased cohorts, there are pairs with positive as well as with negative values of *Δ*_*ij*_. Pairs of nodes for which *Δ*_*ij*_ > 0 corresponds to brain regions which have decreased their normalized shortest communicability path length when Alzheimer disease is present. Those for which *Δ*_*ij*_ < 0 corresponds to brain regions which have increased their normalized shortest communicability path length. A resume of our results for the pairs that increase and decrease the normalized shortest communicability path lengths is given below:

1. Left hemisphere: 31 pairs increased *Ŝ* (*P*_*i*_) and 79 pairs decreased it;
2. Right hemisphere: 6 pairs increased *Ŝ* (*P*_*i*_) and 35 pairs decreased it;
3. Left-Right hemispheres: 55 pairs increased *Ŝ* (*P*_*i*_) and 112 pairs decreased it;
4. Vermis-Left hemisphere: 0 pairs increased *Ŝ* (*P*_*i*_) and 31 pairs decreased it;
5. Vermis-Right hemisphere: 0 pairs increased *Ŝ* (*P*_*i*_) and 50 pairs decreased it.

These results indicate that in total 76.9% of all node pairs which display significant change after Alzheimer disease have decreased the length of the shortest communicability paths connecting them. This is a highly counterintuitive finding because it literally means that the nodes “are closer” to each other in terms of their communicability distance after the Alzheimer disease has appeared. In order to disentangle the meaning of this finding we will consider the example provided in Fig. 5.

We will consider the removal of edges which do not disconnect the graph, which are known as cyclic edges (in the graph in Fig. 5 all edges are cyclic). First we will prove the following result.

Let *Γ* be a graph and let *Γ - e* be the same graph without the cyclic edge *e* = {*a, b*}. Then, if *G*_*pq*_ (*Γ*) = (*e*^*γA*(Γ)^)_*pq*_ we have that 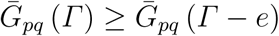, where 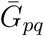 is the average among all pairs of nodes. The proof is given by the fact that if *e* = {*a, b*} is removed, the length of all walks between *a* and *b* will increase, and no other walk will decrease its length. Consequently, the removal of an edge in a graph will drop both the average transmissibility 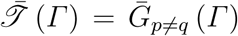 and the average circulability 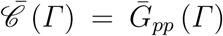 of a disease factor. However, because the communicability distance *ξ*_*pq*_ is the difference between the circulability around the nodes *p* and *q*, and the transmissibility between the two nodes, we have the following situations. Let 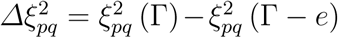, and *ΔG*_*pq*_ = *G*_*pq*_ (Γ) − *G*_*pq*_ (Γ − *e*). Then,

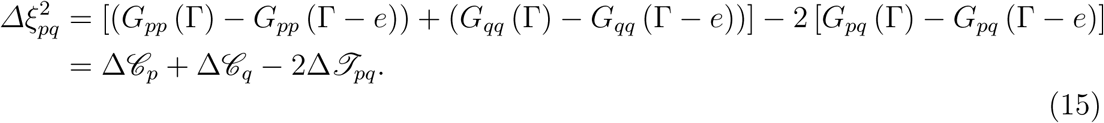

Consequently, when 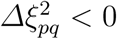 we have that *G*_*pp*_ (Γ) ≲ *G*_*pp*_ (Γ − *e*) and *G*_*qq*_ (Γ) ⪸ *G*_*qq*_ (Γ − *e*), while *G*_*pq*_ (Γ) *⪢ G*_*pq*_ (Γ − *e*). When, 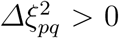 we have that *G*_*pp*_ (Γ) *⪢ G*_*pp*_ (Γ − *e*) and *G*_*qq*_ (Γ) *⪢ G*_*qq*_ (Γ − *e*), while *G*_*pq*_ (Γ) ⪸ *G*_*pq*_ (Γ − *e*).

1. 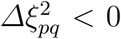 implies that the drop in the transmissibility of the disease factor is bigger than the drop in the circulability around the nodes. That is, in Γ – *e* dominates the circulability to the transmissibility compared to *Γ* ;
2. 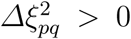 implies that the drop in the circulability of the disease factor is bigger than the drop in the transmissibility around the nodes. That is, in Γ – *e* dominates the transmissibility over the circulability compared to *Γ*.

In Table 1 we report the results that illustrate the previous reasoning for the graph in Fig. 5. First, we report the change in the average shortest path length 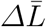 after the removal of the corresponding edges according to the node labelling in Fig. 5. The removal of the edges {1, 2}, {1, 9} and {8, 9} produce significant increase of the communicability shortest path lengths, i.e.,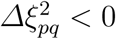. The edge removals {2, 5} and {2, 4} decrease the communicability shortest paths, i.e.,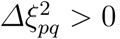. As can be seen for those graphs in which 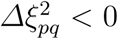 the relative drop of 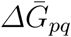, averaged for all edges in the shortest paths, is significantly bigger than that of 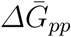, averaged for all nodes. In the case when 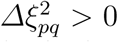, the change in 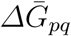 is of the same order than 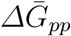. It is reasonably to think that edge removal increases the time needed by the disease factor to infect 100% of the nodes in the network. If we designate this time change by *Δt*_inf_ and calculate it as the minimum time at which all the nodes are infected in the graph using the approximate solution of the SI model (upper bound), we obtain the results given in Table 1. As can be seen when 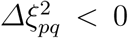 the increase in the infection time is dramatic, ranging from 17 to 30%. In remarkable contrast when 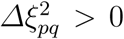 the increase in *t*_inf_ is only 3.8%. In closing, when 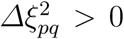 the orifinal network has been transformed into a more efficient one, from the perspective of the transmission of the disease, because in Γ – *e* the circulability around the nodes of the disease factor is sacrified to the transmissibility to other nodes.

**Table 1:** Values of different structural and dynamical parameters for the graph illustrated in Fig. 5 to which edges have been removed. The edges correspond to the labelling of the nodes in the mentioned figure. 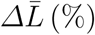 is the percentage of change respect to the original graph in the average communicability shortest path length. 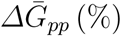 and 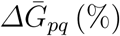 are the percentage of change respect to the original graph for the values of *G*_*pp*_ and *G*_*pq*_ averaged for the nodes and edges in the shortest communicability paths. *Δt*_inf_ (%) is the time needed by a disease factor to infect all the nodes of the corresponding graph in an SI simulation using the approximate solution described below with *β* = 0.005 and initial condition *x*_*i*_(0) = 1*/*9.

| edge | $\Delta\bar{L}$ (%) | $\Delta\bar{G}_{pp}$ (%) | $\Delta\bar{G}_{pq}$ (%) | $\Delta t_{\text{inf}}$ (%) |
| --- | --- | --- | --- | --- |
| $\{1, 2\}$ | -25.4 | 10.0 | 19.2 | 30.5 |
| $\{1, 9\}$ | -23.2 | 15.7 | 26.5 | 25.7 |
| $\{8, 9\}$ | -21.5 | 17.7 | 29.6 | 16.9 |
| $\{3, 5\}$ | -0.04 | 14.9 | 15.6 | 3.8 |
| $\{2, 5\}$ | +0.89 | 15.9 | 15.0 | 3.8 |
| $\{2, 4\}$ | +1.42 | 18.4 | 18.7 | 3.8 |

It is now important to analyze what are the consequences of the decrease in the lengths of the communicability shortest paths that we have observed for Alzheimer diseased brains.The main implication of this observation is that AD produces some damage to the brain, not necessarily only edge removals, which somehow improves the “efficiency” of the resulting networks to propagate the disease in relation to other possible damage scenarios. If we constraint ourselves to the edge-removal scenario and use the previous graph as an example we can conclude that AD has removed those edges which “prioritize” the transmissibility over the circulability of the disease factor, by affecting as least as possible the global rate of contagion in the resulting networks. This is an inference based on the analysis of data produced on AD patients and comparing them with HC. In no case it corresponds to a direct observation of this effect and we call the attention of experimenters to try to falsify this hypothesis. We should remark that in 2013, Tomasi et al. [77] found, using MRI that “a higher degree of connectivity was associated with nonlinear increases in metabolism”. Recent works in network neurosciences have proposed ways to navigate the brain evoiding the “hubs” due to their high energy consumption [78, 79]. Thus, according to our findings, AD produces damages in the connectivity of the brain which drop more significantly the cliquishness–the degree is a first order approach to it–over the transmissibility. Thus, it is also plaussible that propagating the AD through the different regions of an already-damaged brain is less energetic than in the non-damaged one. All-in-all, the damages produced by initial stages of AD seems to improve the capacity of the disease factor to propagate through the network than in the undamaged one.

## 6 Conclusions

There are two main conclusions in the current work. From a theoretical perspective for network neurosciences we have shown that the communicability function–widely used in this field–can be mathematically connected to the solution of a Susceptible-Infected model of disease factor propagation in a brain network. In particular, we have shown that the communicability distance accounts for the difference between the circulability of this disease factor around a brain region (node) and its transmissibility to another region of the brain.

From an application point of view we have provided solid evidences that the shortest communicability path length is significantly better than the shortest topological path length in distinguishing Alzheimer diseased patients from healthy individuals. We have identified a set of 399 pairs of regions for which there are very significant changes in the shortest communicability path length after Alzheimer disease appears, 42% of which interconnect both brain hemispheres and 28% connect regions inside the left hemisphere only. These findings clearly agree with the disconnection syndrome hypothesis of Alzheimer disease. We have also identified 20% of affected regions which connect the vermis with any of the two brain hemispheres. The most significant result is that in 76.9% of these pairs of damaged brain regions there is an increase in the average cliquishness of the intermediate regions which connect them. This results implies that there is a significant increase in energy consumption for communication between these regions in Alzheimer patients in comparison with healthy individuals.

In closing we hope that the current work helps to shed light into important mechanistic aspects of Alzheimer disease as well as in a better understanding of the use of communicability functions for network neuroscience studies.

## 7 Acknowledgments

We wish to thank Robert Smith of the Florey Institute of Neuroscience and Mental Health, Austin, for his valuable suggestions on the image processing pipeline concerning the brain connectivity reconstruction with MRtrix.

Data collection and sharing for this project was funded by the Alzheimer’s Disease Neuroimaging Initiative (ADNI) (National Institutes of Health Grant U01 AG024904) and DOD ADNI (Department of Defense award number W81XWH-12-2-0012). ADNI is funded by the National Institute on Aging, the National Institute of Biomedical Imaging and Bioengineering, and through generous contributions from the following: AbbVie, Alzheimer’s Association; Alzheimer’s Drug Discovery Foundation; Araclon Biotech; BioClinica, Inc.; Biogen; Bristol-Myers Squibb Company; CereSpir, Inc.; Cogstate; Eisai Inc.; Elan Pharmaceuticals, Inc.; Eli Lilly and Company; EuroImmun; F. Hoffmann-La Roche Ltd and its affiliated company Genentech, Inc.; Fujirebio; GE Healthcare; IXICO Ltd.; Janssen Alzheimer Immunotherapy Research and Development, LLC.; Johnson and Johnson Pharmaceutical Research and Development LLC.; Lumosity; Lundbeck; Merck and Co., Inc.; Meso Scale Diagnostics, LLC.; NeuroRx Research; Neurotrack Technologies; Novartis Pharmaceuticals Corporation; Pfizer Inc.; Piramal Imaging; Servier; Takeda Pharmaceutical Company; and Transition Therapeutics. The Canadian Institutes of Health Research is providing funds to support ADNI clinical sites in Canada. Private sector contributions are facilitated by the Foundation for the National Institutes of Health (www.fnih.org). The grantee organization is the Northern California Institute for Research and Education, and the study is coordinated by the Alzheimer’s Therapeutic Research Institute at the University of Southern California. ADNI data are disseminated by the Laboratory for Neuro Imaging at the University of Southern California.

